# Functional and structural features of L2/3 pyramidal cells continuously covary with pial depth in mouse visual cortex

**DOI:** 10.1101/2021.10.13.464276

**Authors:** Simon Weiler, Drago Guggiana Nilo, Tobias Bonhoeffer, Mark Hübener, Tobias Rose, Volker Scheuss

**Author notes:** equal contribution (Simon Weiler, Drago Guggiana Nilo). corresponding author: Volker Scheuss.

## Abstract

Pyramidal cells of neocortical layer 2/3 (L2/3 PyrCs) integrate signals from numerous brain areas and project throughout the neocortex. Within L2/3, PyrCs show functional and structural specializations depending on their pial depth, indicating participation in different functional microcircuits. However, it is unknown whether these depth-dependent differences result from separable L2/3 PyrC subtypes or whether functional and structural features represent a continuum while correlating with pial depth. Here, we assessed the stimulus selectivity, electrophysiological properties, dendritic morphology, and excitatory and inhibitory synaptic connectivity across the depth of L2/3 in the binocular visual cortex (bV1) of female mice. We find that the structure of the apical but not the basal dendritic tree varies with pial depth, which is accompanied by differences in passive but not active electrophysiological properties. PyrCs in lower L2/3 receive increased excitatory and inhibitory input from L4, while upper L2/3 PyrCs receive a larger proportion of intralaminar input. Complementary *in vivo* calcium imaging revealed a systematic change in visual responsiveness, with deeper L2/3 PyrCs showing more robust responses than superficial PyrCs. Furthermore, deeper L2/3 PyrCs are more strongly driven by contralateral than ipsilateral eye stimulation. In contrast, orientation- and direction-selectivity of L2/3 PyrCs are not dependent on pial depth. Importantly, the transitions of the various properties are gradual, and cluster analysis does not support the classification of L2/3 PyrCs into discrete subtypes. These results show that L2/3 PyrCs’ multiple functional and structural properties systematically correlate with their depth within L2/3, forming a continuum rather than representing discrete subtypes.

**SIGNIFICANCE STATEMENT:** Neocortical pyramidal cells in layer 2/3 (L2/3 PyrCs) are crucial for cortical computation and display heterogenous properties. We investigated whether and how these properties vary across the depth of L2/3 and whether L2/3 PyrCs can be subdivided into distinct subtypes. This is important for a better understanding of the coding strategy and information integration processes within L2/3. We find that multiple properties such as morphology, physiology, connectivity, and functional *in vivo* responses of L2/3 PyrCs correlate with cortical depth in mouse visual cortex. These variations are continuous and do not support classification of L2/3 PyrCs into discrete subtypes. In contrast to L5 and L6, PyrCs in L2/3 therefore process information based on a continuous property space.

## Introduction

The mammalian neocortex processes signals in local microcircuits and integrates information from different brain regions across its layers. Excitatory pyramidal cells of layer 2/3 (L2/3 PyrCs) are cortico-cortical projection neurons that exchange information with other neocortical areas. These cells link the main input and output layers of the neocortical circuit (L4, L5/L6) and are therefore a key element in cortical information processing (reviewed in Petersen & Crochet, 2013).

It is well established that neocortical PyrCs are heterogenous with respect to their genetic profiles, morphological and electrophysiological properties, circuit connectivity, and *in vivo* functional response properties (Harris and Shepherd, 2015). In the infragranular layers, several PyrC subtypes have been defined based on specific distinctions in these properties, and such subtypes are thought to form important building blocks for neocortical computations (Vélez-Fort et al., 2014; Kim et al., 2015). This is different in layer 2/3: although PyrCs in L2/3 have been categorized based on single features such as transcriptional profile, morphology or physiology alone, multi-feature clustering has not revealed unambiguous PyrC subtypes in this layer so far (Tasic et al., 2016; Meng et al., 2017; Gouwens et al., 2019; Scala et al., 2021).

This suggests that, rather than originating from discrete, spatially intermingled neuronal subtypes, functional and structural features of L2/3 PyrCs may vary continuously or follow larger scale anatomical gradients, like cortical depth. Indeed, structural, molecular and functional characteristics of L2/3 neurons were found to vary with distance from pia (Kreile et al., 2011; Staiger et al., 2015; Tasic et al., 2016; Gouwens et al., 2019; O’Herron et al., 2020). In mouse visual cortex, individual L2/3 PyrCs are selectively tuned to distinct visual features, such as orientation and direction (Niell and Stryker, 2008; Andermann et al., 2011; Marshel et al., 2011), and continuous depth-dependent changes in these properties have been reported (O’Herron et al., 2020). It was also shown that, similar to other sensory cortical areas (Tasic et al., 2018; Yao et al., 2021), the genetic makeup of PyrCs in the superficial part of L2/3 differs from other L2/3 PyrCs in primary visual cortex (V1) (Tasic et al., 2016), further suggesting that L2/3 is not a functionally homogenous layer. Likewise, morphological and physiological properties are different in the upper compared to the lower part of L2/3 (Gouwens et al., 2019). Additionally, the long-range outputs of L2/3 PyrCs have been shown to vary across L2/3: PyrCs in V1 projecting to specific higher visual areas, such as the anterolateral (AL) or posteromedial (PM) area, reside at different cortical depths (Kim et al., 2020). Interestingly, these cells do not differ in their electrophysiological properties (Kim et al., 2018) and mostly share the same transcriptome (Kim et al., 2020).

Apart from the influence of morphological and electrophysiological characteristics, the visual response properties of L2/3 PyrCs derive from integration of their synaptic inputs within the cortical circuit. Locally, L2/3 PyrCs receive their input through intra-as well as interlaminar excitatory and inhibitory connections, the latter originating from L4 and L5 in mouse V1 (Kätzel et al., 2011; Xu et al., 2016). In particular, interactions between excitatory and inhibitory presynaptic inputs play an important role in shaping the functional response properties of individual L2/3 PyrCs (Rossi et al., 2020). The variance of L2/3 PyrC morphology with pial-depth (Gouwens et al., 2019) together with the fact that different types of inputs target different subcellular compartments (Petreanu et al., 2009) suggests that L2/3 connectivity within the local circuit also depends on pial depth. In rodent somatosensory and auditory cortex such relationship has been observed (Staiger et al., 2015; Meng et al., 2017), where neurons in the superficial compared to the deeper part of L2/3 differ in the amount of input from specific layers and in the horizontal extent from where inputs arise. It remains to be explored whether such depth-dependent variations in intra- and interlaminar connections exist in other sensory cortical areas, and whether these input changes are continuous or discrete within L2/3.

Taken together, it is still unclear whether information is processed by discrete L2/3 PyrC subtypes or by a continuum of neurons with a gradually varying feature set. Furthermore, it remains to be established to which extent L2/3 should be considered a uniform layer and if neuronal properties change with pial depth. Therefore, a systematic approach taking into account multiple structural and functional features of PyrCs across the full extent of L2/3 is needed to better understand the organization of this layer. We therefore assessed how the morpho-electric properties, intra- and interlaminar input connectivity, and visual response properties of excitatory L2/3 neurons are distributed, and how they relate to pial depth. We find that the apical dendritic architecture, the passive intrinsic properties, and the local input sources to L2/3 PyrCs vary systematically with depth. This is accompanied by gradual changes in visual response properties, arguing for a gradually changing microcircuit within L2/3. Finally, the distributions of these features do not support clustering of cells into discrete subtypes, but rather argue for a functional continuum of L2/3 PyrCs.

## Methods

### Animals

All experimental procedures were carried out in compliance with institutional guidelines of the Max Planck Society and the local government (Regierung von Oberbayern). Wild type C57bl/6 female mice (postnatal days P27-P70) were used. Mice were housed under a 12 h light-dark cycle with food and water available ad libitum. *In vitro* brain slice experiments were performed at P30-P70. Craniotomy, virus injections and head plate implantation were performed at P30-P35. *In vivo* imaging was performed at P50-P70. Animals were usually group housed. After cranial window and head plate implantation animals were singly housed. All the experiments were performed during the dark cycle of the animals.

### Solutions

The cutting solution for *in vitro* experiments contained 85 mM NaCl, 75 mM sucrose, 2.5 KCl, 24 mM glucose, 1.25 mM NaH_2_PO_4_, 4 mM MgCl_2_, 0.5 mM CaCl_2_ and 24 mM NaHCO_3_ (310-325 mOsm, bubbled with 95% (vol/vol) O_2_, 5% (vol/vol) CO_2_). Artificial cerebrospinal fluid (ACSF) contained 127 mM NaCl, 2.5 mM KCl, 26 mM NaHCO_3_, 2 mM CaCl_2_, 2 mM MgCl_2_, 1.25 mM NaH_2_PO_4_ and 10 mM glucose (305-315 mOsm, bubbled with 95% (vol/vol) O_2_, 5% (vol/vol) CO_2_). Caesium-based internal solution contained 122 mM CsMeSO_4_, 4 mM MgCl_2_, 10 mM HEPES, 4 mM Na-ATP, 0.4 mM Na-GTP, 3 mM Na-L-ascorbate, 10 mM Na-phosphocreatine, 0.2 mM EGTA, 5 mM QX-314, and 0.03 mM Alexa 594 (pH 7.25, 295-300 mOsm). K-based internal solution contained 126 mM K-gluconate, 4 mM KCl, 10 mM HEPES, 4 mM Mg-ATP, 0.3 mM Na-GTP, 10 mM Na-phosphocreatine, 0.3-0.5% (wt/vol) Neurobiotin tracer and 0.03 mM Alexa 594 (pH 7.25, 295-300 mOsm).

### Acute brain slice preparation

The detailed procedure is described elsewhere (Weiler et al., 2018). Briefly, mice were deeply anesthetized with Isoflurane in a sealed container and rapidly decapitated. Coronal sections of V1 (320 μm, Bregma −1.5 to −3) were cut in ice cold carbogenated cutting solution using a vibratome (VT1200S, Leica). Slices were incubated in cutting solution in a submerged chamber at 34°C for at least 45 min and then transferred to ACSF in a light-shielded submerged chamber at room temperature (21°C) until used for recordings. Brain slices were used for up to 6 hours. A single brain slice was mounted on a *poly-D-lysine coated* coverslip and then transferred to the recording chamber of the microscope while keeping track of the rostro-caudal orientation of the slice.

### Laser Scanning Photostimulation (LSPS)

For uncaging experiments using UV laser light, two different setups were used. Coronal brain slices were visualized with an upright microscope (setup A: BW51X, Olympus; setup B: A-scope, Thorlabs) using infrared Dodt gradient contrast (DGC) with a low magnification UV transmissive objective (4x objective lens) and images were acquired by a high-resolution digital CCD camera. MNI-caged-L-glutamate concentration was 0.2 mM. The bath solution was replaced after 3 h of recording, and bath evaporation was counterbalanced by constantly adding a small amount of distilled H_2_O to the solution reservoir using a perfusor. L2/3 PyrCs in bV1 were targeted using morphological landmarks and then whole cell recordings were performed at high magnification using a 60x objective. Targeted PyrC bodies were at least 50 μm below the slice surface. Borosilicate glass patch pipettes (resistance of 4-5 MΩ) were filled with a Cs-based internal solution for measuring excitatory and inhibitory postsynaptic currents (EPSC: voltage clamp at −70 mV, IPSC: voltage clamp at 0-5 mV). Electrodes also contained 30 μM Alexa 594 for detailed morphological visualization using 2-photon microscopy. Once stable whole-cell recordings were obtained with good access resistance (< 30 MΩ) the microscope objective was switched from 60x to 4x. Mapping experiments were controlled with Ephus software (Suter et al., 2010). The slice was positioned within the CCD camera’s field of view and a stimulus grid (16 x 16 with 69 μm spacing) was aligned to the recorded cell’s soma and the pial surface. Multiple maps were recorded with grid locations stimulated in a pseudo-random fashion (1 ms pulses, 10-15 mW in the specimen plane, 1s interstimulus interval, 2-3 repetitions each with different mapping sequence) for both excitatory and inhibitory inputs.

On setup A, a diode-pumped solid state (DPSS laser Inc.) laser was used to generate 355 nm UV laser pulses for glutamate uncaging. The duration and intensity of the laser pulses were controlled by an electro-optical modulator, a neutral density filter wheel and a mechanical shutter. The laser beam was scanned using voltage-controlled mirror galvanometers. An UV-sensitive photodiode measured the power of the UV laser beam. A dichroic mirror reflected the UV beam into the optical axis of the microscope while transmitting visible light for capturing bright-field images by the CCD camera. The beam passed a tube/scan lens pair in order to underfill the back aperture of the 4x mapping objective resulting in a pencil-shaped beam.

On setup B, the UV laser was an Explorer One 355-1 (Newport Spectra-Physics). The duration and intensity of the laser pulses were directly controlled using analog signals, the built-in software L-Win (Newport Spectra-Physics), a mechanical shutter and neutral density filters. An UV-sensitive photodiode measured the power of the UV laser beam.

Data were acquired with Multiclamp 700 B amplifiers (Axon instruments). Voltage clamp recordings were filtered at 4-8 kHz and digitized at 10 kHz. Data Analysis was performed using custom-written software in MATLAB. The spatial resolution of photostimulation was estimated using excitation profiles (Shepherd and Svoboda, 2005). Excitation profiles describe the spatial resolution of uncaging sites that generate action potentials in stimulated neurons. For this, excitatory as well as inhibitory cells in different layers of bV1 were recorded either in whole-cell or cell-attached configuration using a K-based internal solution in current-clamp mode. Mapping was performed as described above only that the stimulus grid was 8×8 or 8×16 with 50 or 69 μm spacing. The spatial resolution was 60-100 μm depending on cell type and layer (data not shown).

### Intrinsic properties measurements

K-based internal solution was used when recording passive and active electrophysiological properties. Once stable whole-cell recordings were obtained with good access resistance (usually < 30 MΩ) basic electrophysiological properties were examined in current-clamp mode with 1 s long hyper- and depolarizing current injections.

### Image acquisition for morphological imaging

The patch pipette was carefully retracted from the cell after successful recording and filling with Alexa-594. A detailed structural 2-photon image stack of the dendritic morphology of the entire cell was acquired with excitation light of λ=810 nm using ScanImage 4.2 (Pologruto et al., 2003). The structural image stacks typically consisted of 250 sections (1024 x 1024 pixels; 0.3-0.8 μm per pixel) collected in z steps of 1-2 μm.

### Virus dilution, injection and chronic window preparation

The detailed procedure is described elsewhere (Weiler et al., 2018). To co-express the genetically encoded calcium indicator GCaMP6m together with the structural marker mRuby2 (Rose et al., 2016) in a sparse subset of L2/3 neurons, the adeno-associated virus AAV2/1-Syn-FLEX-mRuby2-CSG-P2A-GCaMP6m-WPRE-SV40 (titer: 2.9 x 10^13^ GC per ml, Addgene accession no. 102816) in combination with AAV2/1.CamKII0.4.Cre.SV40 (titer: 1.8 x 10^13^ GC per ml, University of Pennsylvania Vector Core accession no. AV-1-PV2396) were used. The final titer of AAV2/1-Syn-FLEX-mRuby2-CSG-P2A-GCaMP6m-WPRE-SV40 was 1.4 x 10^13^ GC per ml (PBS was used for dilution).

Briefly, surgeries were performed on 32 female C57bl/6 mice that were intraperitoneally (i.p.) anesthetized with a mixture of Fentanyl (0.05 mg kg^-1^), Midazolam (5 mg kg^-1^) and Medetomidine (0.5 mg kg^-1^). Additional analgesic drugs applied were Carprofen (5 mg kg^-1^, subcutaneous, s.c.) before surgery and Lidocaine (10%, topical to skin prior to incision). A section of skin over the right hemisphere starting from the dorsal scalp was removed and the underlying periosteum was carefully removed. A custom-machined aluminum head bar (oval shape, with an 8 mm opening and two screw notches) was carefully placed and angled over the binocular zone of the primary visual area. The precise location of the binocular zone was determined by intrinsic optical signal (IOS) imaging through the intact skull prior to the craniotomy in each animal (see section below). A circular craniotomy (4 mm diameter) centered over the binocular zone of the right primary visual cortex was performed. The premixed virus was injected 200-500 μm below the pial surface at a single site in the binocular zone of V1 (50-100 nl/injection, ~ 10 nl/min ejected by pressure pulses at 0.2 Hz) using glass pipettes and a pressure micro injection system. Additionally, diluted fluorescent retrobeads (1:20 with cortex buffer, Lumafluor Inc.) were pressure injected (10-20 nl/injection, 5 nl/min) medial and lateral to the virus injection site at ~1500 μm from its center. The craniotomy was covered with a glass cover slip and was sealed flush with drops of histoacryl. The head bar and cover glass were then further stabilized by dental cement. After surgery, the animal was injected s.c. with saline (500 μl) and the anesthesia was antagonized by i.p. injection of Naloxone (1.2 mg kg^-1^), Flumazenil (0.5 mg kg^-1^) and Atipamezole (2.5 mg kg^-1^). Carprofen (5 mg kg^-1^, subcutaneous, s.c.) was administered the following two days. *In vivo* imaging was performed not earlier than 2 weeks after virus injection to allow for sufficient indicator expression.

### Intrinsic optical signal imaging

For IOS imaging, the optical axis was orthogonal to the head bar. The brain surface was first illuminated with light of 530 nm to visualize the blood vessel pattern and subsequently with 735 nm for intrinsic imaging in order to localize bV1. Images were acquired using a 4x air objective (NA 0.28, Olympus) and a CCD camera (12 bit, 250×348 pixel, 40 Hz). The camera was focused ~500 μm below the pial surface. Image acquisition and analysis software were custom-written in MATLAB. The visual stimulus was a patch with a size of 20° x 40° displayed randomly to either the left or right eye at two distinct positions next to each other in the central visual field. Within the patch a sinusoidal grating was displayed in eight directions for 7 s (grating direction was changed every 0.6 s) with a temporal frequency of 2 cycles/s and a spatial frequency of 0.04 cycles/degree. Individual trials were separated by 8 s of a full-field gray stimulus (50% contrast). The entire stimulus sequence was applied at least 2 times for each eye and patch position during the surgery before virus injection and at least 3 times at the beginning of the first *in vivo* imaging session

### In vivo 2-photon imaging

L2/3 PyrCs co-expressing GCaMP6m and the bright structural marker mRuby2 (mRuby2-CSG-P2A-GCaMP6m) were imaged *in vivo* using a tunable pulsed femtosecond Ti:Sapphire laser (Newport Spectra-Physics) and a customized commercial 2-photon microscope (16x 0.8 NA water immersion objective; B-Scope I, Thorlabs). The laser was tuned to λ=940 nm in order to simultaneously excite GCaMP6m and mRuby2. After rejecting excitation laser light (FF01-720/25, Semrock), the emitted photons passed through a primary beam splitter (FF560 dichroic, Semrock) and band pass filters (FF02-525/50 and FF01-607/70, Semrock) onto GaAsP photomultiplier tubes (H7422P-40, Hamamtsu) to separate green and red fluorescence.

Multiple imaging planes were acquired by rapidly moving the objective in the z-axis using a high-load piezo z-scanner (P-726, Physik Instrumente). The imaged volume for functional cellular imaging was 250 x 250 x 100 μm^3^ with 4 inclined image planes, each separated by 25 μm in depth. Imaging frames of 512 x 512 pixels (pixel size 0.5 μm) were acquired at 30 Hz by bidirectional scanning of an 8 kHz resonant scanner while beam turnarounds were blanked with an electro-optic modulator (Pockels cell). Imaging was performed between 130-400 μm below the pial surface. Excitation power was scaled exponentially (exponential length constant ~150 μm) with depth to compensate for light scattering in tissue with increasing imaging depth. The average power for imaging was <50 mW, measured after the objective. The optical axis was adjusted orthogonal to the cranial window. ScanImage 4.2 (Pologruto et al., 2003) and custom written hardware drivers were used to control the microscope.

After functional characterization of L2/3 PyrCs, at least two high-resolution structural image stacks with different field of view sizes were acquired at λ=940 nm/1040 nm. 1) 450 sections (512 x 512 pixels) with a pixel size of 0.5 μm collected in z-steps of 1.4 μm (imaged volume of 256 x 256 x 630 μm^3^); 2) 350 sections (512 x 512 pixels) with a pixel size of 1.9 μm collected in z-steps of 2 μm (imaged volume of 972 x 972 x 700 μm^3^).

Experiments were performed under light anesthesia. Data acquisition started ~45 min after an i.p. injection of Fentanyl (0.035 mg kg^-1^), Midazolam (3.5 mg kg^-1^) and Medetomidine (0.35 mg kg^-1^). Additional doses of anesthetics (25% of induction level) were subcutaneously injected every 45-60 mins to maintain the level of anesthesia. Ophthalmic ointment was applied to protect the eyes. Mice were fixed under the microscope by screwing the metal head-plate to two posts. Stable thermal homeostasis was maintained by using a heated blanket throughout the imaging session. Eye and pupil positions were recorded with two cameras (DMK 22BUC03, The Imaging Source Europe GmbH) throughout *in vivo* imaging.

### Visual stimulation

Visual stimuli were generated using the MATLAB Psychophysics Toolbox extension and displayed on a gamma-corrected LCD monitor ((Brainard, 1997), http://psychtoolbox.org). The screen measured 24.9 x 44.3 cm, had a refresh rate of 60 Hz and was positioned in portrait orientation 13 cm in front of the eyes of the mouse, providing a viewing angle of ~45 deg on each side from the center of the monitor. The monitor was adjusted in position (horizontal rotation and vertical tilt) for each mouse to align with the horizontal visual axis and to cover the binocular visual field (−15° to 35° elevation and −25° to 25 azimuth relative to midline). The presented stimulus area was chosen to subtend binocular visual space and the rest of the screen was uniformly grey (50% contrast). An OpenGL shader was applied to all presented stimuli to correct for the increasing eccentricity on a flat screen relative to the spherical mouse visual space (Marshel et al., 2011). Randomly alternating monocular stimulation of the eyes was achieved by motorized eye shutters and custom MATLAB scripts.

For all visual stimuli presented, the backlight of the LED screen was synchronized to the resonant scanner, switching on only during the bidirectional scan turnaround periods when imaging data were not recorded (Leinweber et al., 2014). The mean luminance with 16 kHz pulsed backlight was 0.01 cd/m^2^ for black and 4.1 cd/m^2^ for white.

To measure visually evoked responses, the right or left eye was visually stimulated in random order using drifting black and white square wave gratings of eight directions with a temporal frequency of 3 cycles/s and a spatial frequency of 0.04 cycles/degree. Stimulation duration for moving gratings was 5 s interleaved by 6 s of a full-field grey screen. Trials were repeated 4 times per eye and direction.

### Morphological reconstruction and analysis

The reconstruction of dendritic cell morphology was performed manually using the Simple Neurite Tracer of ImageJ (Schindelin et al., 2012). Reconstructions were quantitatively analyzed in MATLAB and with the open-source TREES toolbox (Cuntz et al., 2011). The radial distance was measured as the Euclidean distance from the soma to each segment terminal. The total length was measured as the sum of all internode sections’ lengths of the neurite. For Sholl analysis, the number of intersections between dendrites and concentric spheres centered on the soma was determined at increasing distances from the soma (20 μm increments). The distance to peak branching was measured as the distance of maximal dendritic branching from the soma. The width/height ratio was measured as the overall maximum horizontal extent divided by the overall maximum vertical extent.

### Intrinsic properties extraction

Electrophysiological parameters were extracted using the PANDORA Toolbox (Günay et al., 2009) and custom-written software in MATLAB. The active, single spike parameters were measured using the first spike evoked by current injection (at Rheobase). The parameters were measured/calculated and defined in the following way:

Passive:

1. Resting membrane potential (V_rest_): The membrane potential measured after break-in.
2. Membrane time constant, *τ*_m_ (ms): This was estimated using an exponential fit to the recovery of the voltage response following hyperpolarizing step currents.
3. Input resistance, R_IN_ (MΩ): Estimated by the linear fit of the I-ΔV curve (using subthreshold de- and hyperpolarizing pulses (−30:10:30 pA).
4. Sag in percentage (Sag ratio): 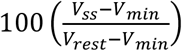 where V_ss_ is the voltage at steady-state, V_rest_ the resting membrane potential and V_min_ the minimum voltage reached during hyperpolarizing current injections of −300 pA.
5. Rheobase (pA): The minimum current amplitude of infinite duration required for action potential generation. Measured by depolarizing current pulses (10:10:300 pA).

Active:

1. Minimal membrane voltage during Afterhyperpolarization (APV_min_): This was estimated as the membrane potential minimum during the period of the AHP.
2. Peak membrane voltage of action potential (APV_peak_).
3. Threshold voltage at action potential initiation (APV_thresh_).
4. The maximal slope of the action potential (APV_slope_): The maximal rate of rise of membrane voltage during the spike rise phase.
5. Membrane voltage at action potential half-height (APV_half_).
6. Amplitude of the action potential (APV_amp_): Amplitude calculated as difference between the voltage at APV_thresh_ and APV_peak_.
7. Maximal amplitude of AHP (AHP): It was measured as the difference between the APV_thresh_ and APV_min_.
8. Spike frequency, APfreq_max_ (Hz): The maximum action potential number evoked by step-current injections divided by the pulse duration. Measured at the depolarizing current pulse, that evoked maximum action potential number (10-400 pA).

### Input map analysis

The spatial resolution of LSPS by UV glutamate uncaging was calculated based on the size of the excitation profiles as the mean weighted distance from the soma (d_soma_) of AP generating stimulation sites using the following equation (Shepherd and Svoboda, 2005):

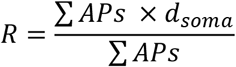

LSPS by UV glutamate uncaging induces two types of responses (Shepherd and Svoboda, 2005): 1) Direct glutamate uncaging responses originating from direct activation of the glutamate receptors on the recorded neuron by uncaged glutamate. 2) Synaptic responses originating from the activation of synaptic glutamate receptors on the recorded neuron by glutamate release from presynaptic neurons stimulated by LSPS. Responses to the LSPS stimulation protocol (both for EPSCs and IPSCs) were quantified in the 150 ms window following the uncaging light-pulse, since this is the time window where evoked activity is observed in most cases. Considering the diversity of responses encountered in these experiments, a heuristic analysis scheme was devised to address the main observed cases:

1. Traces without response were excluded by only considering those responses with a deflection higher than 2 S.D. over the baseline at any point. Additionally, traces that only had a significant response in one repetition were also excluded.
2. Then, purely synaptic responses, i.e. those that correspond only to activation of the presynaptic neuron via uncaged glutamate were selected by taking the traces that passed the 2 S.D. threshold only after a 7 ms window from the offset of stimulation.
3. For responses that did not pass the previous criterion, inspection by eye indicated that several of them presented all the identifiable features of purely synaptic responses but seemed to cross the threshold slightly earlier than 7 ms. An additional set of experiments performed on a subset of cells, where maps were measured before and after application of TTX (and hence before and after only direct responses were present) were performed to characterize these intermediate cases (~5% of the total number of traces). These experiments showed that by using a secondary window of 3.5 ms, the average contribution of a direct response to the overall response in these intermediate traces is ~20 % (data not shown). Therefore, this secondary window was used to include a second batch of traces into the synaptic response pool.
4. Finally, those traces that did not pass the secondary window were then blanked, and a 4dimensional interpolation method (MATLAB function “griddatan”) was used to infer their temporal profiles based on their 8 neighboring pixel activities in space and time. In the TTX experiments (data not shown) every position with a direct response was observed to have a synaptic component, but the summation of this synaptic component and the overlapping direct component is non-linear. Therefore, this interpolation method was used to extract the synaptic component partially masked in the raw traces by the direct response. The approach relies on the observation that the synaptic responses of neighboring positions are similar across time, therefore indicating that information on the synaptic responses masked by direct responses is contained in the responses surrounding them. These interpolated responses were then incorporated into the maps as synaptic responses. For excitatory input maps, the first two stimulation rows were excluded since L1 contains no excitatory neurons (Jiang et al., 2015) and excitatory input from L1 originated from cells in L2/3-L5 having apical tuft dendrites in L1, which fired action potentials in exceptional cases when their tufts were stimulated in L1 (Dantzker and Callaway, 2000).

For Principal Component Analysis (PCA) on input maps, the input maps were aligned based on the soma position of each cell. This involved shifting the maps vertically an integer number of stimulus rows until all the somata were in the same row. Subsequently, all maps were normalized and used as features for PCA. The combined excitation-inhibition PCA decomposition was then calculated. For this, the feature vectors from excitation and inhibition for each map were concatenated, yielding a 512 element feature vector that was then used for the decomposition. The first three principal component weights for each input map were extracted (carrying roughly 60% of the variance in the dataset).

The data includes input maps of 70 L2/3 PyrCs from a previously obtained data set (Weiler et al., 2020).

### UMAP embedding

Uniform Manifold Approximation and Projection was utilized to visualize the distribution of different properties across the data on a cell by cell basis. The computational details of UMAP are described elsewhere (McInnes et al., 2018). Briefly, UMAP embeds data points from a high dimensional space into a 2D space preserving their high dimensional distances in a neighborhood. This permits effective visualization of the connections between data points. A UMAP implementation in MATLAB developed by Meehan, Meehan and Moore (https://www.mathworks.com/matlabcentral/fileexchange/71902) was utilized. The respective principal components for morphology, electrophysiology, input maps and *in vivo* functional responses were used as the embedding parameters. The number of neighbors was 15 and the minimum distance was 0.1 (default parameters). The embedded points were color-coded depending on the normalized pial-depth.

### In vivo imaging analysis

Custom-written MATLAB code was used for image and data analysis. For IOS imaging analysis, the acquired images were high-pass filtered and clipped (1.5%) to calculate blank-corrected image averages for each condition. Additionally, a threshold criterion (image background mean + 4 x standard deviation) was set to determine the responsive region within the averaged image. The mean background value of the non-responsive region was subtracted from each pixel and all pixel values within the responsive area were summed to obtain an integrated measure of response strength.

In the case of 2-photon calcium imaging, the use of GCaMP6m in combination with mRuby2 gave the possibility to perform ratiometric imaging (Rose et al., 2016). Image sequences were full-frame corrected for tangential drift and small movements caused by heart beat and breathing. An average of 160 image frames acquired without laser excitation was subtracted from all frames of the individual recording to correct for PMT dark current as well as residual light from the stimulus screen. Cell body detection was based on the average morphological image derived from the structural channel (mRuby2) for each recording session. ROIs (region of interest) were drawn manually and annotated. The fluorescence time course was calculated by averaging all pixel values within the ROI on both background-corrected channels, followed by low-pass filtering (0.8 Hz cut-off) and by subtraction of the time-variable component of the neuropil signal (pixel average within a band of 15 μm width, 2 μm away from the ROI circumference, excluding overlap with other selected cells and neuropil bands, neuropil factor r of 0.7 (Kerlin et al., 2010)). The green and red fluorescence signal were estimated as:

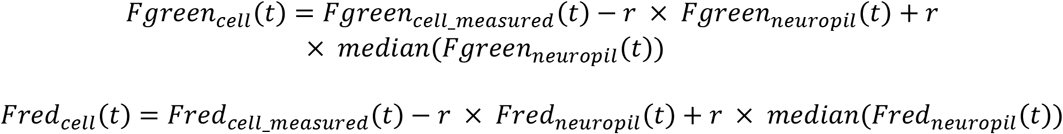

The ratio R(t) was then calculated as:

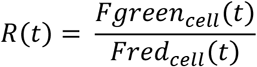

Residual trends were removed by subtracting the 8^th^ percentile of a moving 14 s temporal window from R(t). ΔR/R_0_ was calculated as:

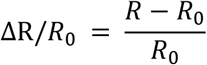

where R_0_ is the median of the mean baseline fluorescence ratios over a 1 s period preceding visual stimulation in each trial. Visual responses were quantified as mean fluorescence ratio change over the full stimulus interval both in individual trials and the trial-averaged mean fluorescence ratio.

Visual responsiveness was tested with a one-way ANOVA performed over all trials with and without visual stimulus. Neurons with *p*-values < 0.05 were identified as visually responsive. OD was determined by the OD index (ODI):

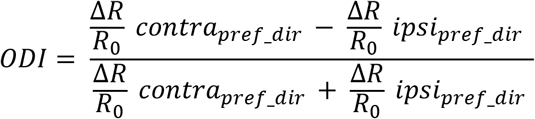

Where an ODI value of 1 or −1 indicates exclusive contra- and ipsilateral dominance, respectively.

Global orientation selectivity index (gOSI) was computed as 1 − circular Variance (circ. Var.):

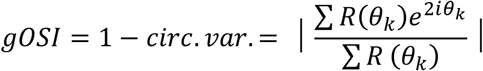

and global direction selectivity index (gDSI) was computed as:

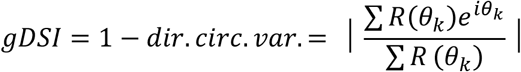

*R*(*θ_k_*) is here the mean response to the direction angle (*θ_k_*) (Mazurek). Perfect orientation and direction selectivity is indicated with gOSI and gDSI of 1, whereas a gOSI and gDSI value of 0 indicates no orientation or direction selectivity, respectively. The preferred orientation and direction as well as tuning width were computed by fitting a double-Gaussian tuning curve to the responses as previously described (Carandini and Ferster, 2000). The tuning width was extracted as the sigma of the fitted curve. The goodness-of-fit was assessed by calculating R^2^ and only cells with R^2^ >0.3 were included in the analysis.

For binocular cells, the preferred orientation was defined as the one from the dominant eye, as determined by the sign of the ODI.

### Statistics

Data are reported as mean ± standard error of the mean (SEM). Correlation coefficients were calculated as Pearson’s correlation coefficient. Before comparison of data, individual data sets were checked for normality using the Kolmogorov-Smirnov Goodness-of-Fit test. None of the data sets considered in this study was found to be normally distributed. Therefore, paired or unpaired nonparametric statistics (Wilcoxon rank sum test) were used for comparison. Twotailed tests were used unless otherwise stated. Correction of multiple comparison was performed by the Benjamini & Hochberg procedure (Benjamini and Hochberg, 1995). Asterisks indicate significance values as follows: *p<0.05, ** p<0.01, *** p<0.001.

## Results

### Morphological properties of L2/3 pyramidal cells vary gradually with pial depth

The dendritic architecture of a cell constrains the sampling of potential synaptic inputs and thereby controls information integration. To study the variations of dendritic architecture across L2/3, 189 Alexa 594-filled L2/3 PyrCs in mouse bV1 were manually reconstructed (36 of these were included from a previously collected data set (Weiler et al., 2020)).

Three representative examples of dendritic morphologies across L2/3 are shown in Fig. 1A. The data set covers the whole cortical depth of L2/3, with cells reconstructed in upper as well as lower parts of the layer (Fig. 1A, B). Given that apical and basal dendrites are targeted by different types of inputs (feedback vs. feedforward, (Petreanu et al., 2009)), we separately characterized the apical and basal dendritic architecture by Sholl analysis (Fig. 1C). In addition, we extracted sets of commonly used morphological parameters for the apical and basal dendrites (Table 1). Overall, the parameters are either related to dendritic length (e.g., total length, maximal horizontal extent, distance to peak Sholl crossing, see Methods) or to dendritic complexity (e.g., number of branch points, peak number of Sholl crossings, see Methods).

**Figure 1:**
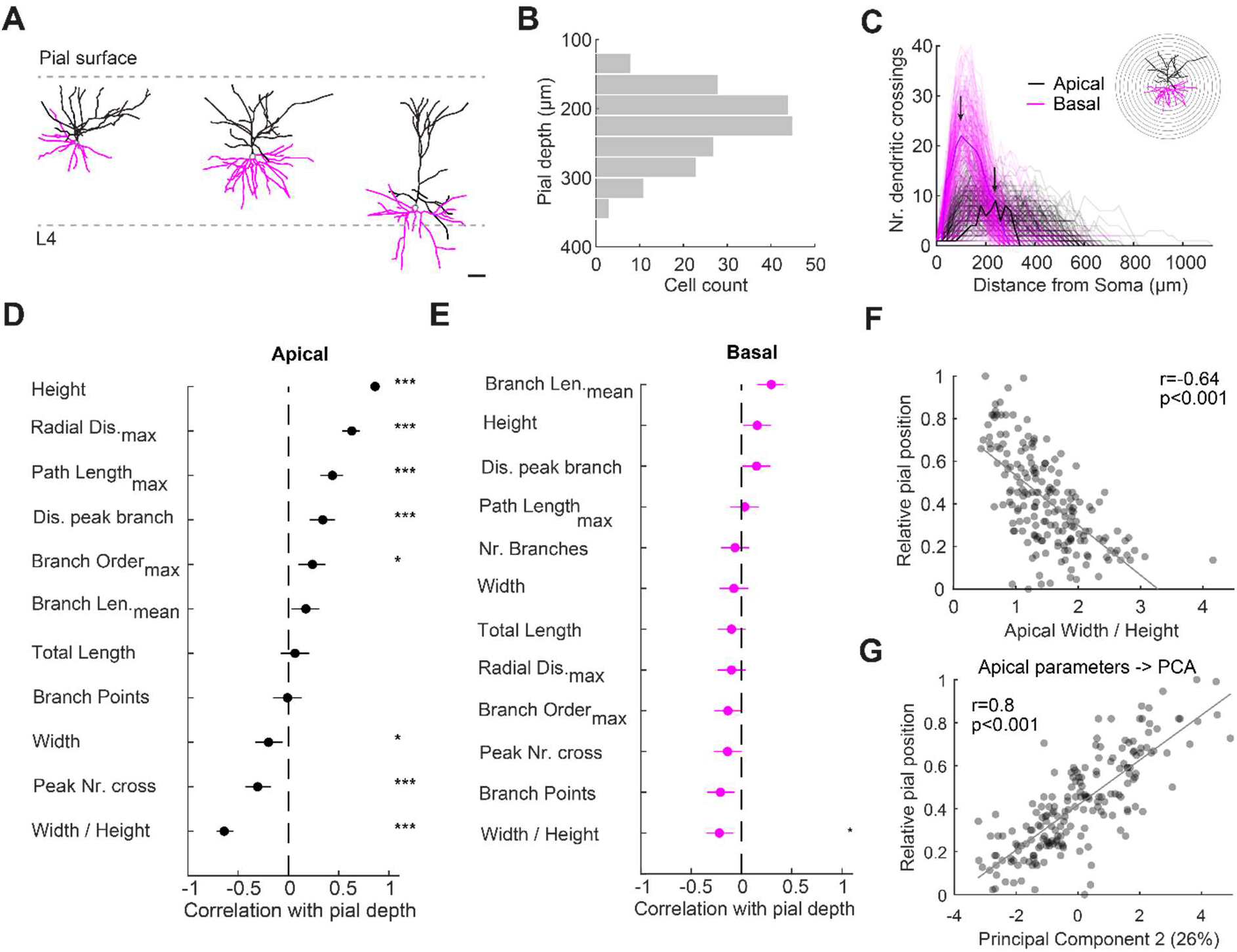
Apical but not basal dendritic morphology of L2/3 PyrCs changes with pial depth. **A** Reconstructed dendritic morphology of PyrCs in the upper, middle, and lower part of L2/3 (scale bar: 50 μm). Apical dendrites, black, basal dendrites, magenta. **B** Distribution of distances to the pial surface of morphologically reconstructed neurons within L2/3. **C** Sholl analysis for apical and basal dendrites. The number of crossings was determined using concentric spheres centered around the soma with 20 μm increments. Bold lines refer to the example cell in inset. Arrows indicate the peak number of crossings for the example cell. **D** Correlations between apical dendritic tree parameters and pial depth sorted in descending order. Error bars are 95% confidence intervals. Asterisks indicate significant correlations. Multiple comparison corrected using Benjamini & Hochberg procedure with false discovery rate (FDR) of 0.05 (Benjamini and Hochberg, 1995). **E** Same as D for basal dendrite parameters. **F** Relative soma position within L2/3 (0 − top, 1 - bottom of L2/3) plotted against ratio of width over height of the apical tree. Linear fit is indicated in grey. Pearson correlation coefficient r indicated at top right. **G** Relative soma position within L2/3 plotted against principal component 2 weight for apical tree morphology. Percentage indicates variance explained by this principal component. Linear fit is indicated in grey. All data presented is from n=189 cells, from 76 mice.

**Table 1.** List of parameters used for morphological analysis of apical and basal dendritic trees with their corresponding average values and contributions to the first three principal components from principal component analysis performed separately for apical and basal tree (eigenvalues PC1-PC3, n=189 cells, from 76 mice).

| # | Description | Mean ± SEM | PC1 | PC2 | PC3 |
| --- | --- | --- | --- | --- | --- |
| **Apical dendrite** |  |  |  |  |  |
| 1 | Radial Dis.<sub>max</sub>: Maximal radial distance from soma | 223.94 ± 3.47 μm | 0.28 | 0.47 | 0.23 |
| 2 | Total length: Total length of tree | 2015.1 ± 46.28 μm | 0.36 | -0.08 | 0.39 |
| 3 | Path Length<sub>max</sub>: Maximal path length from soma | 173.6 ± 3.99 μm | 0.07 | -0.08 | 0.21 |
| 4 | Branch Points: Number of branch points | 16.4 ± 0.4 | -0.29 | -0.2 | 0.31 |
| 5 | Branch Order<sub>max</sub>: Maximal branch order | 7.98 ± 0.15 | 0.49 | -0.25 | -0.1 |
| 6 | Branch Length<sub>mean</sub>: Mean branch length | 59.6 ± 0.68 μm | -0.35 | 0.12 | 0.29 |
| 7 | Width / Height: Width / Height of tree | 1.5 ± 0.04 | 0 | 0.06 | 0.73 |
| 8 | Width: maximal horizontal span | 288.91 ± 6.18 μm | 0.32 | -0.32 | 0.14 |
| 9 | Height: maximal vertical span | 217.72 ± 4.18 μm | -0.44 | -0.19 | 0.05 |
| 10 | Peak Nr. Cross: Peak number of crossing (Sholl Analysis) | 11.09 ± 0.26 | -0.21 | -0.2 | 0.03 |
| 11 | Dis. Peak branch: Distance to peak crossing (Sholl Analysis) | 199.69 ± 7.93 μm | -0.02 | 0.68 | -0.06 |
| **Basal dendrite** |  |  |  |  |  |
| 1 | Radial Dis.<sub>max</sub>: Maximal radial distance from soma | 141.06 ± 3.14 μm | 0.34 | 0.46 | 0.15 |
| 2 | Total length: Total length of tree | 2394 ± 57.37 μm | 0.27 | -0.13 | 0.44 |
| 3 | Path Length<sub>max</sub>: Maximal path length from soma | 157.04 ± 7.99 μm | 0.13 | -0.04 | -0.14 |
| 4 | Branch Points: Number of branch points | 23.36 ± 0.6 | -0.33 | -0.01 | 0.41 |
| 5 | Branch Order<sub>max</sub>: Maximal branch order | 8.33 ± 0.19 | -0.28 | 0.13 | 0.01 |
| 6 | Branch Length<sub>mean</sub>: Mean branch length | 49.87 ± 0.57 μm | 0.28 | -0.28 | 0.02 |
| 7 | Width / Height: Width / Height of tree | 1.26 ± 0.02 | 0.14 | 0.02 | -0.36 |
| 8 | Width: maximal horizontal span | 240.22 ± 4.47 μm | 0.23 | 0.01 | 0.66 |
| 9 | Height: maximal vertical span | 197.19 ± 3.21 μm | 0.18 | -0.44 | -0.11 |
| 10 | NB: Number of basal trees | 5.86 ± 0.1 μm | 0.65 | 0.07 | -0.14 |
| 11 | Peak Nr. Cross: Peak number of crossing (Sholl Analysis) | 21.7 ± 0.54 | 0.05 | 0.08 | -0.05 |
| 12 | Dis. Peak branch: Distance to peak crossing (Sholl Analysis) | 125.6 ± 4.14 μm | 0.03 | 0.69 | -0.05 |

To compare depth-dependent changes, we sorted the apical and basal dendritic tree parameters according to their correlation with the cell’s depth within L2/3 in descending order (Fig. 1D, E, Pearson’s correlation coefficient). This showed that most (8 out of 11) apical tree parameters were significantly correlated with pial depth. In contrast, only 1 out of 12 basal tree parameters was significantly correlated with pial depth. Most prominently, the apical trees of neurons located in the more superficial part of L2/3 had the largest horizontal extent (width). Since the apical dendrites of all cells reached the pial surface, we also observed a strong relation between vertical extent (height) and pial depth (Fig. 1D, F). To eliminate potential redundancies in the information carried by these parameters, we performed principal component analysis separately for the apical and basal dendrites. For the apical dendrite, the first three principal components, explaining approximately 75% of variance, were significantly correlated with pial depth (PC1: r=0.15, p<0.05; PC2: r=0.8, p<0.001; PC3: r= −0.29, p<0.001, Pearson’s correlation coefficient) with PC2 showing the strongest correlation (Fig 1G, see Table 1 for principal components eigenvalues). For the basal dendrite, principal component 1 and 2, but not principal component 3 were uncorrelated with pial depth (PC1: r=-0.11, p=0.14; PC2: r=0.12, p=0.1; PC3: r= −0.32, p<0.001, Pearson’s correlation coefficient).

Taken together, the apical dendritic architecture of L2/3 PyrCs systematically varies with pial depth, whereas the basal tree morphology does not.

### Passive but not active electrical properties of L2/3 pyramidal cells vary gradually with pial depth

Besides the dendritic architecture, the intrinsic electrical characteristics influence the functional properties of neurons. To determine the electrophysiological properties of PyrCs across the depth of L2/3, we analyzed the responses of 137 L2/3 PyrCs to hyper- and depolarizing somatic current injections (Fig. 2A, B). We measured five passive and eight active intrinsic properties (Table 2). Again, we sorted the passive and active properties according to their correlation with the cell’s depth within L2/3 in descending order (Fig. 2C, D). While four out of five passive intrinsic properties significantly correlated with cortical depth, none of the eight active properties did. Specifically, more superficial L2/3 PyrCs had a larger input resistance (R_IN_) and at the same time slower membrane time constants (*τ*_m_) compared to PyrCs in the lower part of L2/3 (Fig. 2E). Performing PCA on the passive intrinsic properties also showed a correlation between pial depth and the first two principal components, explaining approximately 75% of variance (Fig. 2F, PC1 vs. pial depth: r=-0.32, p<0.001, PC2 vs. pial depth: r=-0.18, p<0.05, Pearson’s correlation coefficient, see Table 2 for principal components eigenvalues).

**Figure 2:**
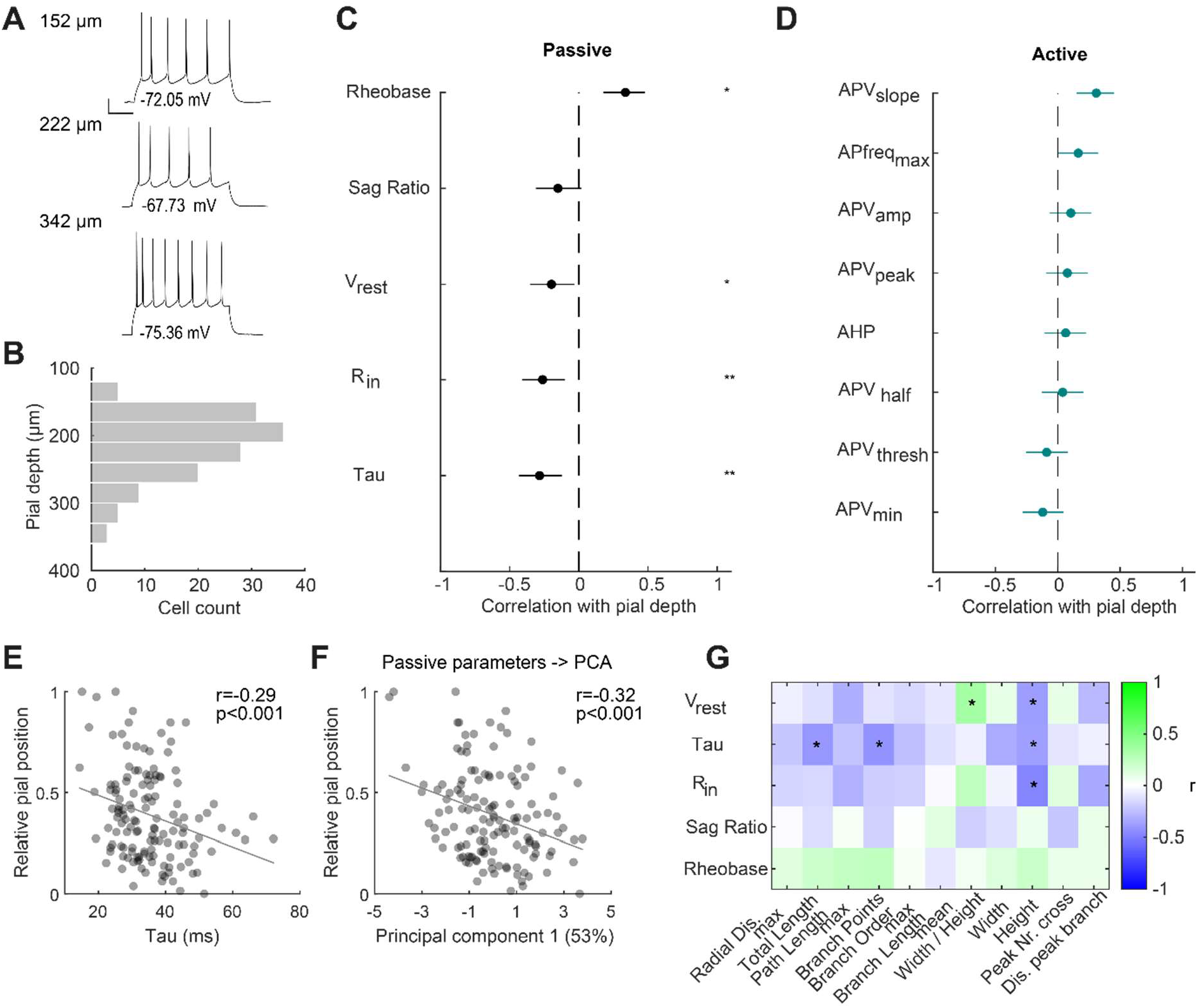
Passive but not active electrophysiological properties change with pial depth. **A** Voltage response to a depolarizing step current (Rheobase +30 pA) of three representative L2/3 PyrCs at increasing pial depth (scale bars: 10 mV, 10 ms). **B** Distribution of distances to the pial surface of electrophysiologically characterized neurons within L2/3 (n=137, from 41 mice). **C** Correlations between passive intrinsic properties and pial depth sorted in descending order. Error bars are 95% confidence intervals. Asterisks indicate significant correlations. Multiple comparison corrected using Benjamini & Hochberg procedure with FDR of 0.05. **D** Same as C for active electrophysiological properties. **E** Relative soma position within L2/3 plotted against membrane time constant. Linear fit is indicated in grey. Pearson correlation coefficient r indicated at top right. **F** Relative soma position within L2/3 plotted against PC1 weight for passive intrinsic properties. Percentage indicates variance explained by this principal component. **G** Correlations between morphological parameters for apical tree and passive intrinsic properties (n=32 cells, from 14 mice). Color indicates the Pearson correlation coefficient between the pair of parameters according to the color bar on the right. Asterisks indicate significant correlations.

**Table 2.**
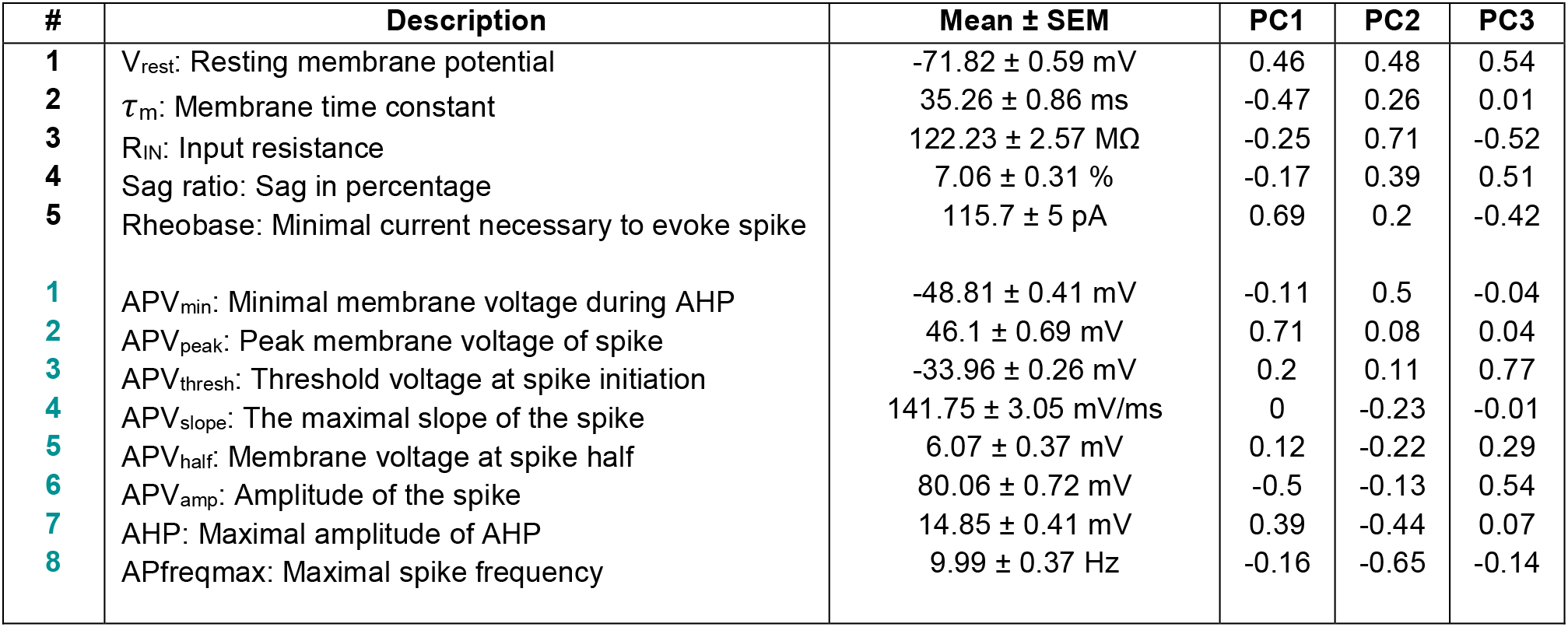
The 5 passive (black) and 8 active (green) extracted electrophysiological parameters with their corresponding average values and contributions to the first three principal components from principal component analysis performed separately for passive and active electrophysiological parameters (eigenvalues PC1-PC3, n=137 cells from 41 mice).

| # | Description | Mean ± SEM | PC1 | PC2 | PC3 |
| --- | --- | --- | --- | --- | --- |
| 1 | $V_{rest}$: Resting membrane potential | $-71.82 \pm 0.59$ mV | 0.46 | 0.48 | 0.54 |
| 2 | $\tau_m$: Membrane time constant | $35.26 \pm 0.86$ ms | -0.47 | 0.26 | 0.01 |
| 3 | $R_{IN}$: Input resistance | $122.23 \pm 2.57$ MΩ | -0.25 | 0.71 | -0.52 |
| 4 | Sag ratio: Sag in percentage | $7.06 \pm 0.31$ % | -0.17 | 0.39 | 0.51 |
| 5 | Rheobase: Minimal current necessary to evoke spike | $115.7 \pm 5$ pA | 0.69 | 0.2 | -0.42 |
| 1 | APV<sub>min</sub>: Minimal membrane voltage during AHP | $-48.81 \pm 0.41$ mV | -0.11 | 0.5 | -0.04 |
| 2 | APV<sub>peak</sub>: Peak membrane voltage of spike | $46.1 \pm 0.69$ mV | 0.71 | 0.08 | 0.04 |
| 3 | APV<sub>thresh</sub>: Threshold voltage at spike initiation | $-33.96 \pm 0.26$ mV | 0.2 | 0.11 | 0.77 |
| 4 | APV<sub>slope</sub>: The maximal slope of the spike | $141.75 \pm 3.05$ mV/ms | 0 | -0.23 | -0.01 |
| 5 | APV<sub>half</sub>: Membrane voltage at spike half | $6.07 \pm 0.37$ mV | 0.12 | -0.22 | 0.29 |
| 6 | APV<sub>amp</sub>: Amplitude of the spike | $80.06 \pm 0.72$ mV | -0.5 | -0.13 | 0.54 |
| 7 | AHP: Maximal amplitude of AHP | $14.85 \pm 0.41$ mV | 0.39 | -0.44 | 0.07 |
| 8 | APfreqmax: Maximal spike frequency | $9.99 \pm 0.37$ Hz | -0.16 | -0.65 | -0.14 |

In a subset of L2/3 PyrCs, we obtained both the dendritic morphology as well as the intrinsic properties. In line with the above-described depth-dependent changes of apical tree and passive properties, we observed that some features covaried in this subset of cells. We found that the total length as well as the complexity of the apical tree was negatively correlated with τm, confirming that the dendritic structure influences the passive properties, as has been previously demonstrated ((Bekkers and Häusser, 2007) but see (Deitcher et al., 2017)).

In summary, several passive but no active electrical properties of L2/3 PyrCs systematically vary with pial depth.

### Spatial connectivity of L2/3 pyramidal cells varies with pial depth

Given the functional response heterogeneity of L2/3 PyrCs in V1 (Niell and Stryker, 2008; Andermann et al., 2011; Marshel et al., 2011), and the aforementioned changes in morpho-electric properties with pial depth, we wondered whether the excitatory and inhibitory microcircuits, in which L2/3 PyrCs are embedded, also systematically vary based on the cell’s position in L2/3. We therefore mapped the monosynaptic intra- and interlaminar excitatory and inhibitory inputs to 147 L2/3 PyrCs via UV-glutamate uncaging in acute coronal brain slices of bV1 (Callaway and Katz, 1993; Dantzker and Callaway, 2000). We recorded excitatory and inhibitory input in the same cells, and thus were able to assess their relationship on a cell-by-cell basis across the depth of L2/3.

We observed that input maps varied in the laminar and horizontal distribution of synaptic input sources depending on the postsynaptic cell location within L2/3 (Fig. 3A, B). For quantification, we peak-normalized the input maps, computed the input fractions per row and column of the stimulus grid, and sorted these based to their correlation with the cell’s depth within L2/3 in descending order (Fig. 3C, D). As reported for auditory cortex (Meng et al., 2017), we observed that the fraction of excitatory and inhibitory input from L4 was positively correlated with the distance between the cell and the pia (Fig. 3C, E, r=0.41 and r=0.3, p<0.001, Pearson’s correlation coefficient) with more superficial cells receiving less fractional excitation and inhibition from L4 in comparison to deeper cells. Excitatory input from L2/3 displayed the opposite correlation (Fig. 3C, r=-0.3, p<0.001, Pearson’s correlation coefficient). Such correlation was not present for L5 inputs and inhibitory input from L2/3 (Fig. 3C, L5 EX, r=-0.04, p=0.65; L5 IN, r=0.17, p=0.09; L2/3 IN, r=0, p=0.98, Pearson’s correlation coefficient). Since most PyrCs received stronger excitation than inhibition from L4 regardless of their location, the difference between excitation and inhibition did not significantly correlate with pial depth (Fig. 3E, bottom).

**Figure 3:**
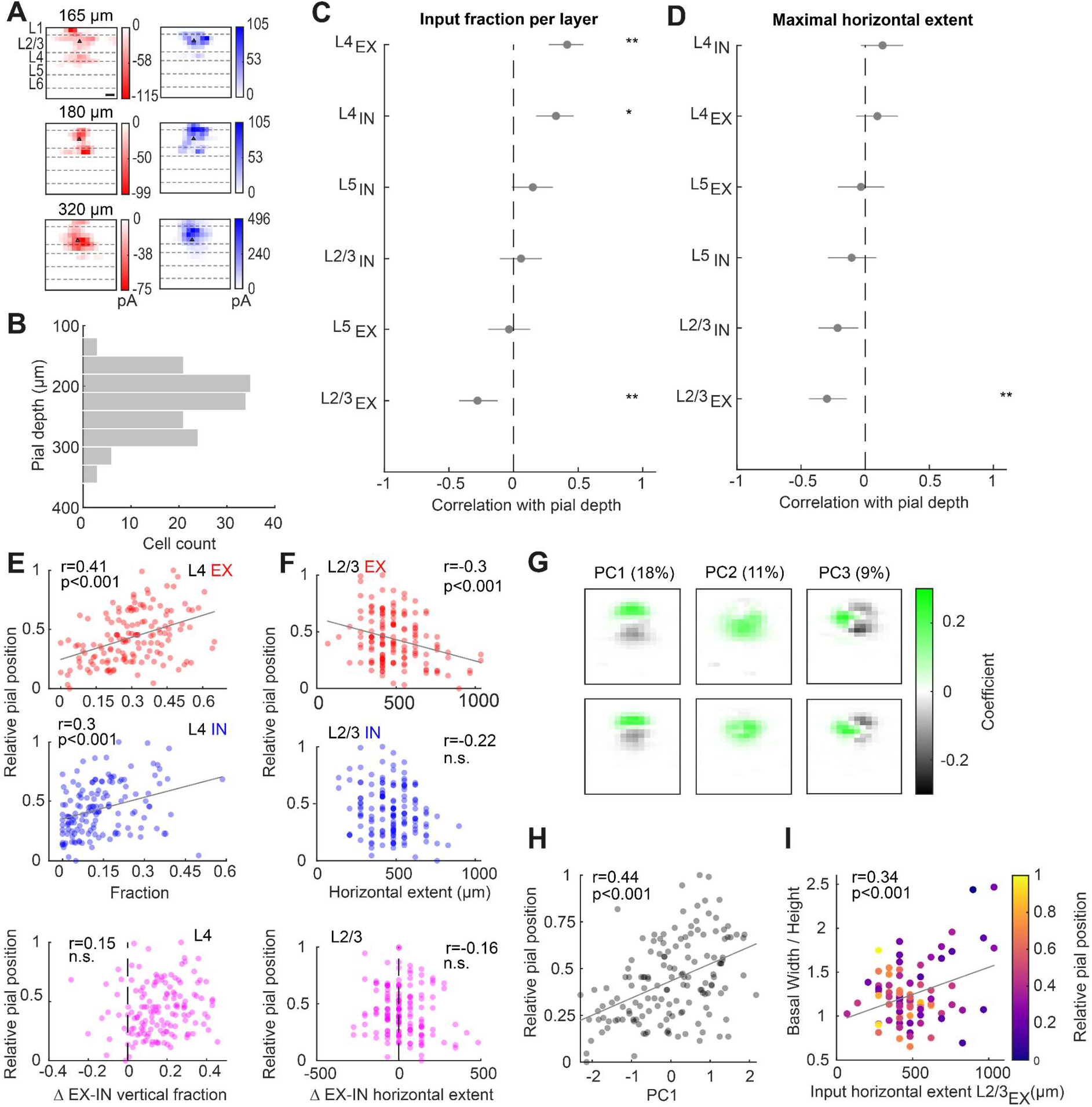
Functional intra- and interlaminar excitatory and inhibitory input connectivity changes with pial depth. **A** Representative, peak normalized excitatory (red) and inhibitory (blue) input maps for PyrCs in the upper, middle, and lower part of L2/3 (scale bar 100 μm). **B** Distribution of distances to the pial surface of functionally mapped neurons within L2/3 (n=147, from 56 mice). **C** Correlations between excitatory and inhibitory input fractions per layer and pial depth sorted in descending order. Error bars are 95% confidence intervals. Asterisks indicate significant correlations. Multiple comparison corrected using Benjamini & Hochberg procedure, FDR=0.05. **D** Same as C for the maximal horizontal extent of input from each layer. **E** Relative pial depth plotted against excitatory (top) and inhibitory (middle) input fractions arising from L4 as well as difference of both (bottom) (n=147 cells, from 56 mice). Pearson correlation coefficient r indicated at top of each plot. Linear fit is indicated in grey. **F** Same as E for maximal horizontal extent of excitatory and inhibitory input from L2/3. **G** Input maps of the first three principal component eigenvalues. Principal component analysis (PCA) using the combined 16×16 normalized excitatory and inhibitory input maps. Before performing PCA, the input maps were vertically and horizontally aligned (see Methods). Explained variance for each principal component is indicated at top**. H** Pial depth plotted against PC1 weight (n=147 cells, from 56 mice). Linear fit is indicated in grey. **I** Ratio of width over height of the basal tree plotted against maximal horizontal extent of excitatory input from L2/3. Color indicates relative soma position within L2/3 according to the color bar on the right (n=97 cells, from 47 mice). For E, F, H and I, the Pearson correlation coefficient r is indicated at the top.

Along the horizontal axis, the maximum spatial extent of the excitatory but not the inhibitory input distributions in L2/3 are negatively correlated with pial depth, with cells displaying a larger spatial extent in upper compared to lower L2/3 (Fig. 3F). This suggests that the extent of cortical space across which L2/3 PyrCs integrate within-layer information increases gradually with decreasing pial depth.

To account for potential redundancies in the information carried by the measured parameters, PCA was performed on the entire set of 16×16 pixel input maps, at the same time for excitation and inhibition (Fig. 3G, see Methods). Prior to PCA, the input maps were horizontally and vertically aligned based on the soma position of each cell. The input maps corresponding to the first three principal components (“eigenmaps”, Fig. 3G) explained ~60% of the variance for both excitatory and inhibitory inputs. Importantly, the first and the third principal component significantly correlated with the pial depth, even though we accounted for cell location information by alignment before performing PCA (Fig. 3H, PC1 vs. pial depth, r=0.44, p<0.001; PC3 vs. pial depth, r=-0.23, p<0.01, Pearson’s correlation coefficient). This indicates that the input pattern itself contains information about the cell location. The principal components were strongly related to the vertical and horizontal spatial features of the input maps described above. For example, while the PC1 weight was significantly correlated with the difference between the excitatory and inhibitory input fraction in L2/3 (r=0.35, p<0.001, Pearson’s correlation coefficient), the PC3 weight was significantly correlated with the difference between the excitatory and inhibitory input fraction in L4 (r=-0.35, p<0.001, Pearson’s correlation coefficient).

Finally, in a subset of 97 L2/3 PyrCs, we reconstructed the dendritic morphology and mapped the functional input in the same cells, enabling us to directly compare the covariations of these two qualities. We observed numerous correlations between the functional input features and the apical dendritic parameters given their respective pial depth dependencies. We therefore focused on the relation between basal tree architecture - which does not covary with depth (Fig. 1) - and spatial input arrangement since most local input terminates on the basal tree (Shepherd et al., 2005; Petreanu et al., 2009). We found that the ratio between the width and height of the basal tree positively correlated with the horizontal extent of the functional input in L2/3 (Fig. 3I, r=0.34, p<0.01, Pearson’s correlation coefficient). Additionally, these parameters were the only ones that significantly correlated with pial depth when considering the basal tree and the horizontal presynaptic input (see also section above). This suggests that, as pial depth decreases, L2/3 PyrCs gradually sample more widely distributed functional input across cortical space. This larger input sampling is potentially achieved via horizontally extended basal trees.

Taken together, these results show that L2/3 PyrCs display a gradual change in the spatial organization of their input distributions with pial depth.

### *In vivo* L2/3 pyramidal cells show depth-dependent variations in stimulus response amplitude and ocular dominance, but not in tuning heterogeneity

How do the observed gradual changes in the different properties relate to visual responses of L2/3 PyrCs in bV1 *in vivo?* Previous recordings in L2/3 of mouse monocular V1 showed a gradual change in overall responsiveness and orientation as well as direction selectivity with pial depth (O’Herron et al., 2020). However, the depth-dependent distribution of other features like eye-specific responsiveness have remained unaddressed so far.

To better understand eye-specific responsiveness, feature selectivity as well as the change of binocularity across the depth of L2/3 in bV1, we performed *in vivo* 2-photon calcium imaging (Fig. 4A). For this, we expressed GCaMP6m in L2/3 PyrCs (Weiler et al., 2018), and imaged across depths ranging from 150-400 μm (Fig. 4A, B). We extracted the following visually evoked response features for each cell: Preferred orientation and direction, global orientation and direction selectivity index (gOSI, gDSI), tuning width, maximum response amplitude at the preferred stimulus direction and ocular dominance. To quantify ocular dominance, we computed the ocular dominance index (ODI; ranging from −1 to 1, with ODI<0 indicating ipsilateral and ODI>0 indicating contralateral dominance, Fig. 4B). To better compare depth-dependent changes, we sorted response features in descending order according to their correlation with the cell’s depth within L2/3 (Fig. 4C).

**Figure 4:**
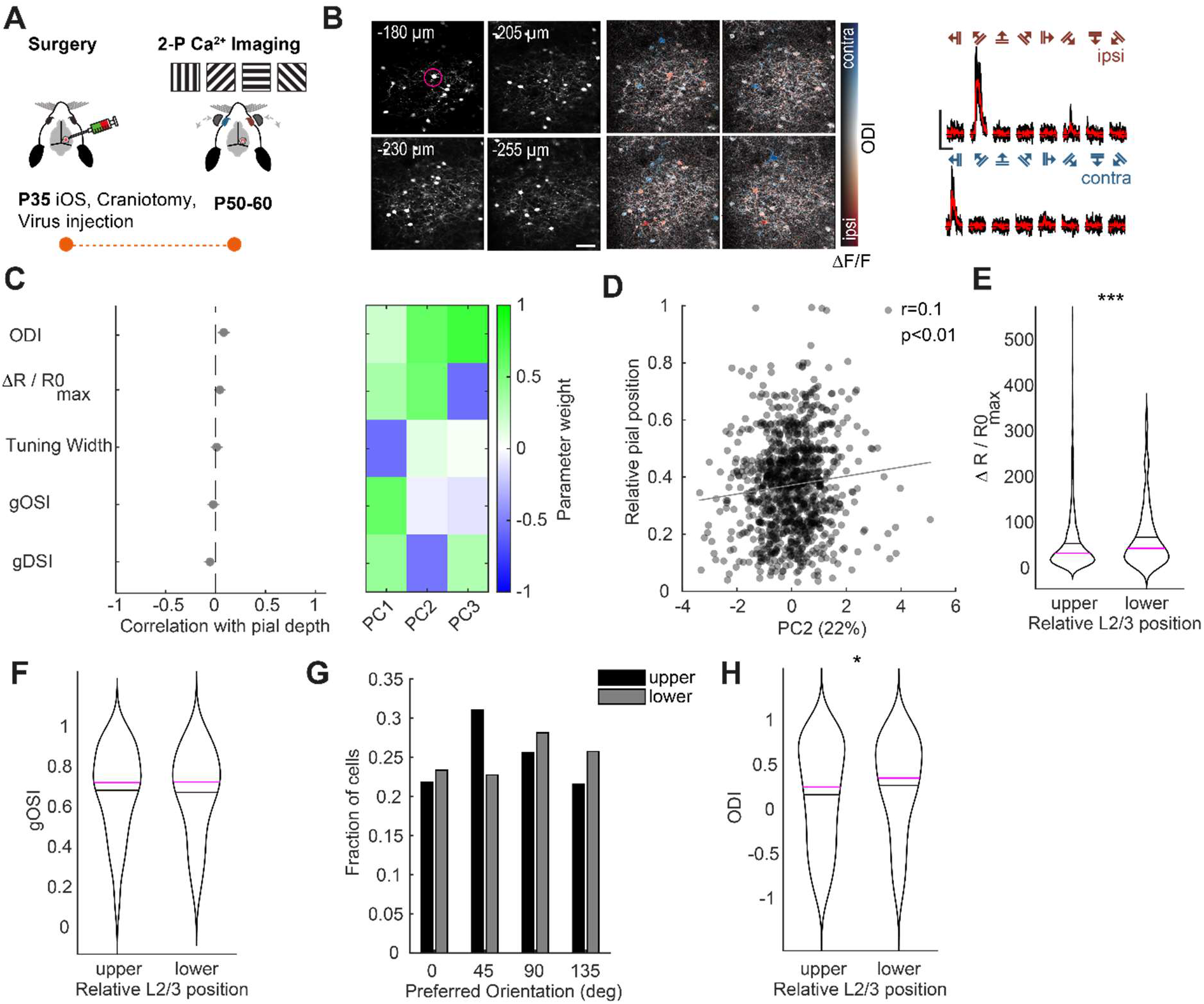
*In vivo* response amplitude and ocular dominance are different between upper and lower L2/3 PyrCs in binocular V1. **A** Experimental pipeline for *in vivo* 2-photon calcium imaging experiments: Binocular visual cortex was identified through the skull by using intrinsic optical signal imaging (iOS, see (Weiler et al., 2018)). Viral injections were then placed into bV1 and a cranial window implanted. After 2-3 weeks of viral expression, moving gratings of different orientations and directions were displayed in front of the mouse. Shutters allow for independent stimulation of either eye. **B** Example image volumes for one animal (four slices acquired with image plane depth increment of 25 μm, scale bar: 50 μm). Left, structural channel: frame-averaged mRuby2 fluorescence. Middle, color-coded response map of individual L2/3 PyrCs. Red and blue hues indicate ipsilateral (ODI<0) and contralateral dominance (ODI>0), respectively. Right, calcium transients of an example neuron in response to ipsi- or contralateral eye stimulation (scale bars: △R/R_0_=200%, 10 s). **C** Left, correlations between visually evoked response features (global orientation and direction selectivity index, gOSI and gDSI, respectively; ocular dominance index, ODI; maximal response to preferred orientation, R/R0_max_) and pial depth sorted in descending order. Error bars are 95% confidence intervals (gOSI, gDSI, tuning width: n=1216 cells, 32 mice; ODI, R/R0_max_: n=1103 cells, from 32 mice). Multiple comparison corrected using Benjamini & Hochberg procedure, FDR=0.05. Right, contributions of the five visually evoked response parameters to the first three principal components. **D** Pial depth plotted against PC2 weight. Pearson correlation coefficient r indicated at top left. Linear fit is indicated in grey (n=1021 cells, from 32 mice). **E** Violin plots of maximal response amplitude for upper and lower L2/3 PyrCs in bV1. Black line indicates mean, magenta line indicates median (n=908 cells for upper, n=226 cells for lower part, from 32 mice). Asterisks indicate significant difference. **F** Same as E for the global orientation selectivity index (gOSI, n=908, n=226 cells, from 32 mice). **G** Distribution of preferred orientation for upper and lower L2/3 PyrCs. **H** Violin plots of ocular dominance index (ODI) for upper and lower L2/3 PyrCs. Black line indicates mean, magenta line indicates median (n=908, n=226 cells, from 32 mice). Asterisks indicate significant difference.

Although some response features displayed correlations with pial depth, these were not significant (after correction for multiple comparison) and far smaller than the correlations observed with morphological, electrophysiological and input map parameters. However, performing PCA on the *in vivo* response features yielded a significant correlation between pial depth and the second principal component (Fig. 4C, D, PC2 vs. pial depth: r=0.1, p<0.01, Pearson’s correlation coefficient). Moreover, when dividing the data into two halves based on the relative pial depth, we observed further depth-dependent differences: PyrCs in the lower part of L2/3 showed significantly larger visually evoked responses compared to PyrCs in the upper part (Fig. 4E, p<0.001, Wilcoxon rank-sum). Importantly, the overall proportion of visually responsive PyrCs was similar across the depth of L2/3 (upper half: 51%, lower half: 47% of all structurally detected PyrCs, see Methods).

Given the previously described depth-dependent changes of orientation selectivity within monocular V1 (O’Herron et al., 2020), we next compared the global orientation selectivity index (gOSI) across the depth of L2/3 (see Methods). The gOSI was similar for PyrCs in the upper and lower part of L2/3, both when including all cells (Fig. 4F) or only cells with strong preferred response amplitude (third quartile, c.f. O’Herron, data not shown). Similarly, the preferred orientations of orientation selective cells (gOSI>0.25) were equally represented in the upper or lower part of L2/3, although there was a slightly higher fraction of PyrCs preferring more oblique oriented gratings (45 degrees) in the superficial part of the layer (Fig. 4G).

When comparing the ocular dominance of PyrCs across the depth of L2/3, we found a gradual change in eye dominance, with cells in the lower part displaying on average significantly larger contralateral eye dominance (Fig. 4H, p<0.05, Wilcoxon rank-sum). This suggests that eye dominance is differentially distributed throughout L2/3.

In summary, in addition to gradual changes of morpho-electric properties and functional input connectivity, several *in vivo* stimulus response properties of L2/3 PyrCs in bV1 also change with pial depth.

### No evidence for distinct subtypes of L2/3 pyramidal cells based on structural and functional properties

We describe depth-dependent changes in several properties that have been used to categorize PyrCs into subtypes in the past (Vélez-Fort et al., 2014; Kim et al., 2015; Gouwens et al., 2019). Consequently, we next wondered whether these variations across L2/3 justify the classification of PyrCs into discrete subtypes.

For evaluating the presence of clusters in the different data sets, we used the extracted principal components followed by a Dip test (Hartigan, 1985; Adolfsson et al., 2019) to asses multimodality in the principal component weights (see Methods). We found that the weights of the first three principal components for morphology, intrinsic properties, spatial distribution of functional input as well as visually evoked response features did not show significant multimodality, arguing against the presence of distinct clusters (Fig. 5A, Hartigan’s Dip test). Moreover, when plotting first and second PC weights against each other, no clear separation was observed for any of the properties (Fig. 5B). This holds also for basal dendritic tree morphology and active electrophysiological properties that do not show correlations with pial depth (data not shown). This suggests that even though L2/3 PyrCs display quantitative differences in their various properties, these differences do not justify the separation of L2/3 PyrCs into discrete subpopulations of cells.

**Figure 5:**
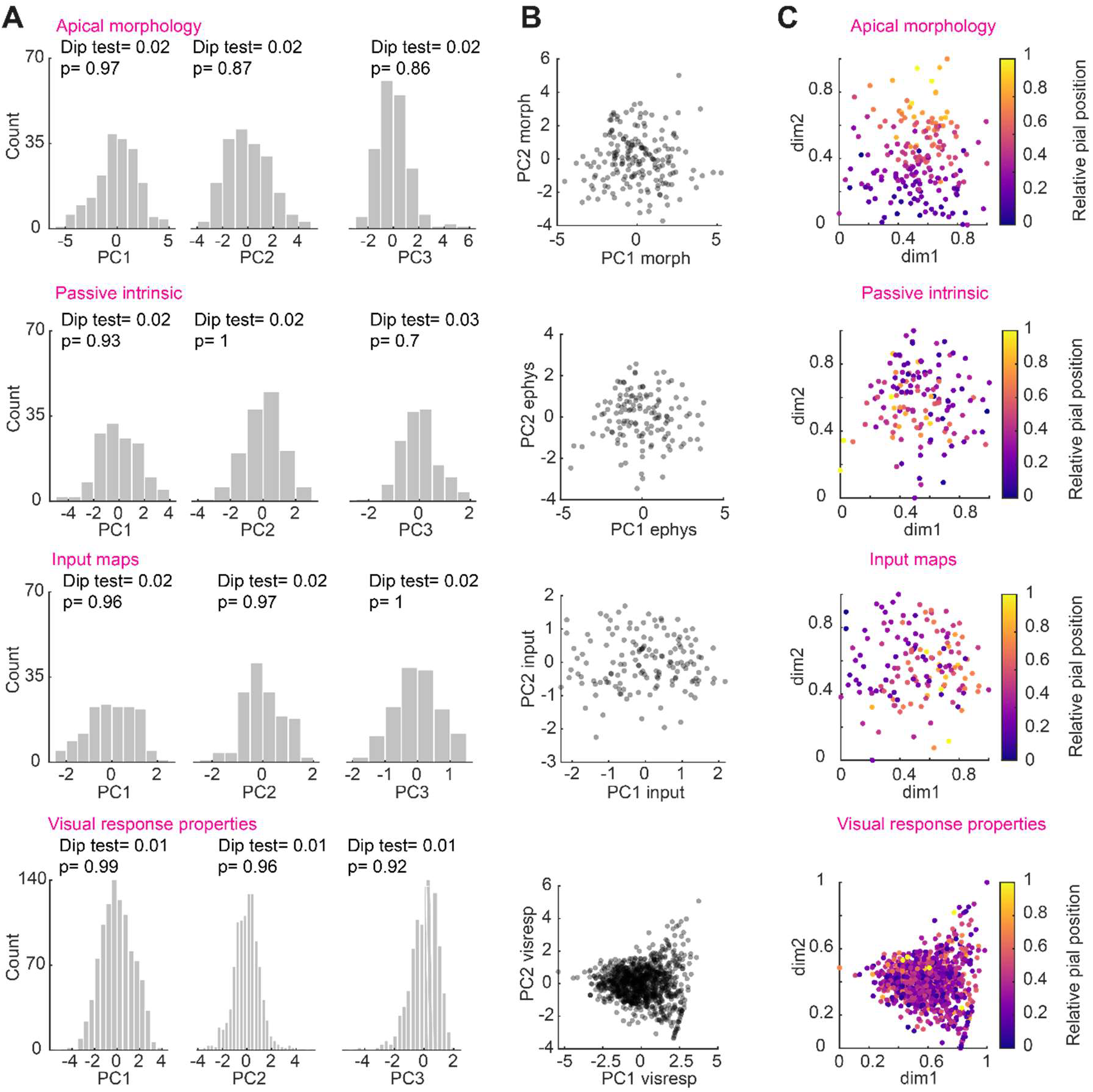
Continuum-like variation of dendritic morphology, passive electrophysiological properties, functional input and visually evoked response properties with pial depth. **A** Distribution of principal component weights and Dip test results for multimodality for the first three principal components calculated for apical tree morphology, passive intrinsic properties, functional excitatory and inhibitory input maps and visually evoked response properties (from top to bottom). **B** Principal component weights PC1 and PC2 from A plotted against each other. **C** UMAP projections color-coded for relative pial position. The UMAP embedding was performed using the first three principal component weights of the respective data sets. Dimension 1 (dm1) and 2 (dm2) are plotted. Data sets from top to bottom: n=189 cells, from 76 mice; n=137 cells, from 41 mice; n=147 cells, from 56 mice; n=1021cells, from 32 mice.

Alternatively, rather than forming separate clusters, L2/3 PyrCs appear to form a single, but inhomogeneous set of neurons whose properties follow a depth-dependent continuum. To illustrate this better, we displayed individual cells in two-dimensional UMAP (Uniform Manifold Approximation and Projection) plots for the different data sets (Fig. 5C), via an embedding based on the first three principal components in each case. The data points aggregated together in single quasi-continuous clouds, rather than separating into well-delineated clusters. However, by color-coding cells according to their pial depth in the UMAP plots, gradients become visible that show how morphology, electrophysiological properties, and input maps systematically vary with pial depth.

In conclusion, morpho-electric features, local excitatory and inhibitory inputs as well as visually evoked response properties of L2/3 PyrCs continuously vary across the depth of visual cortex but this variability does not indicate clusters.

## Discussion

Our study shows that PyrCs vary in multiple properties across the vertical extent of L2/3: 1) The apical dendritic tree progressively spans less horizontal but more vertical space with increasing depth. 2) Passive but not active intrinsic properties gradually change with pial depth. 3) PyrCs in the lower part of L2/3 receive stronger ascending input from L4 compared to PyrCs in the upper part, whereas the horizontal extent of excitatory input is larger for upper vs. lower L2/3 PyrCs. 4) Visual response properties such as ocular dominance and response amplitude show depth-dependent changes. All these changes take place continuously and, thereby, do not justify categorization of L2/3 PyrCs into discrete subtypes.

### Gradually changing morpho-electric properties of L2/3 pyramidal cells

When considering the architecture of their apical tree, PyrCs displayed a morphological continuum across L2/3. PyrCs in lower L2/3 had a long apical dendrite with a tuft, whereas PyrCs in upper L2/3 showed shorter but wider apical trees that branched profusely in L1, as previously described in monocular V1 (Larkman and Mason, 1990; Gouwens et al., 2019) and other sensory cortical areas (Staiger et al., 2015). Interestingly, the total length as well as the number of branch points of the apical tree did not significantly vary between PyrCs located in the upper or lower part of L2/3, similar to other sensory cortical areas (Staiger et al., 2015). Hence, PyrCs throughout L2/3 could in principle sample a comparable number of synaptic inputs, although they display variations in their horizontal as well as vertical extent.

In contrast to the apical tree, the basal dendritic trees did not show any strong relation with pial depth in the present study. This is in line with previous reports showing that basal dendritic trees do not significantly vary across sensory cortical layers (Bielza et al., 2014; Kanari et al., 2019).

The morphological architecture of apical dendrites has been shown to be associated with specific active electrophysiological properties, such as firing patterns (Mainen and Sejnowski, 1996; Deitcher et al., 2017), or passive properties, such as input resistance (Tyler et al., 2015). Numerous studies have reported differences in passive electrophysiological properties of superficial vs. deep L2/3 PyrCs (Zaitsev et al., 2012; Staiger et al., 2015; Van Aerde and Feldmeyer, 2015). The most prominent and consistent difference is that more superficial L2/3 PyrCs show a higher input resistance as well as a slower membrane time constant compared to lower L2/3 PyrCs ((Staiger et al., 2015; Van Aerde and Feldmeyer, 2015; Luo et al., 2017), but see (Deitcher et al., 2017)). Similarly, we found a significant negative correlation of input resistance and membrane time constant with cortical depth in L2/3 PyrCs of mouse bV1.

Additionally, analyzing correlations between morphology and electrophysiology directly in the same cells, we found that total dendritic length and dendritic complexity are negatively correlated with the membrane time constant. The input resistance variance resulted in differences in neuron excitability (as measured via Rheobase in our study). Therefore, cells in the upper regions of L2/3 could in principle be more strongly activated with the same input strength compared to lower L2/3 cells. Indeed, L2 PyrCs in monkey V1 show higher levels of ongoing activity compared to L3 PyrCs (Gur and Snodderly, 2008).

Taken together, the gradual depth-dependent changes in morpho-electric properties of L2/3 PyrCs shape the input and output relationship of these neurons, and ultimately influence the functional information processing across this layer.

### Depth-dependent laminar circuits and functional response properties of L2/3 pyramidal cells

Following the depth-dependent morpho-electric variations of L2/3 PyrCs, we found that the spatial organization of excitatory and inhibitory intracortical inputs to L2/3 gradually changes with cortical depth. A depth-dependent change of intracortical connectivity in L2/3 was also observed in primary somatosensory as well as primary auditory cortex using a similar circuit mapping approach (Staiger et al., 2015; Meng et al., 2017). These studies found that L2/3 PyrCs close to the L4 border receive more ascending excitatory L4 input compared to L2/3 PyrCs close to the L1 border, consistent with our results. Moreover, superficial L2/3 PyrCs received stronger intralaminar excitatory input compared to PyrCs closer to L4. Additionally, the excitatory horizontal extent of input coming from L2/3 was greater for cells in the upper part compared to cells in the lower part of L2/3, similar to the auditory cortex (Meng et al., 2017). However, in visual cortex we only observed this for inputs from within L2/3 and not from any other layer, in contrast to the auditory cortex.

The gradual change of input sources reported here suggests a functional continuum: L2/3 PyrCs at the border to L4 predominately receive ascending feedforward input from L4 in conjunction with L4-mediated inhibition. The contribution of L4 input becomes progressively smaller in the superficial part, where ultimately intralaminar input dominates.

We found that the visually evoked response amplitude was larger in lower L2/3 PyrCs compared to more superficial L2/3 PyrCs, in line with a recent report in monocular V1 (O’Herron et al., 2020). Strong L4 input paired with direct thalamic input (Morgenstern et al., 2016) to PyrCs in the lower part of L2/3, could lead to a stronger feedforward drive compared to upper L2/3 PyrCs, and thereby to the observed differences in response amplitudes. Other *in vivo* tuning properties, such as orientation selectivity, were not significantly different across the depth of L2/3 in our study. This is at odds with previous studies in the monocular part of V1 (Gur and Snodderly, 2008; O’Herron et al., 2020), where the orientation selectivity was stronger in superficial L2/3. Future studies need to address whether this discrepancy is due to a difference in the depth-dependent distribution of this particular property in L2/3 between the monocular and binocular visual cortex, or whether the difference arises from different types of visual stimulation (full field visual stimulation vs. centered stimulation covering only binocular visual space; 1.5 Hz vs. 3 Hz temporal frequency).

With respect to ocular dominance, we find that L2/3 PyrCs closer to the border to L4 are on average dominated by the contralateral eye. This degree of contralateral dominance could in principle be inherited from L4 and/or direct thalamocortical projections (Morgenstern et al., 2016) but future research would be needed to address this contralateral bias.

Taken together, depending on where PyrCs and their corresponding input sources are located, the functional connectivity may directly influence specific functional response properties.

### Absence of well-defined clusters of L2/3 pyramidal cells

The observed depth-dependent variations in the different types of properties extracted in the present study did not support clustering due to their unimodal distributions which argues against subdivision of L2/3 PyrCs into discrete cell types. Likewise, also adding all parameters of the respective data sets that were uncorrelated with pial depth, lead to unimodal distributions and therefore did not support clustering. We thus did not find discrete subtypes of L2/3 PyrCs, which is different from the auditory cortex, where clustering was demonstrated on laminar input fractions, however, without prior testing for multimodality (Meng et al., 2017). Instead, we find a continuum of cellular properties across this layer (Scala et al., 2021). It would be of interest to apply the presented clusterability tests (Adolfsson et al., 2019) on different data sets for L2/3 PyrCs from other cortical regions, both in rodents as well as other species, to test the generalizability of a depth-dependent functional continuum within L2/3 across cortical areas.

Why is it that there is a continuum-like parameter distribution of the different properties within L2/3? One reason could be the associative role of L2/3 in comparison to other layers. For example, an important output route of information from L2/3 PyrCs is via L5 and L6 PyrCs. In contrast to L2/3 PyrCs, L5 and L6 PyrCs separate into distinct subtypes based on the same parameters investigated in this study (Vélez-Fort et al., 2014; Kim et al., 2015; Tasic et al., 2016; Gouwens et al., 2019). The most crucial differences between the infragranular layers and L2/3 are their output projections and their computational role. L5 and L6 contain intratelencephalic (IT) as well as extratelencephalic (ET) neurons, whereas L2/3 only contains IT neurons (Harris and Shepherd, 2015; Peng et al., 2021). Furthermore, PyrCs in L2/3 employ a different coding scheme compared to the infragranular layers. L2/3 PyrCs use sparse coding, whereas PyrCs in infragranular operate with a dense coding scheme (reviewed in (Harris and Mrsic-Flogel, 2013; Petersen and Crochet, 2013)). This indicates that computations in L5 and L6 are performed with projection-specific divisions, whereas within L2/3, such divisions in “hardware” are largely absent, with individual neurons being rather embedded in different IT (cortical-cortical) subcircuits, serving the associative role of this layer.

In summary, numerous neuronal properties of PyrCs gradually change with cortical depth in L2/3. This makes L2/3 a unique cortical layer, where information processing is based on pyramidal neurons with a continuous property space rather than discrete neuronal subtypes.

## Acknowledgements

We are grateful to Volker Staiger for cell tracing as well as technical support, to Michael Myoga for helping to build one *in vitro* setup and to Pieter Goltstein for software. This study was supported by the Max Planck Society and the German Research Foundation (DFG, the Collaborative Research Center SFB870_A08, reference number 118803580; V.S. and M.H.).

## Data availability statement

The datasets generated and analyzed during the current study are available from the corresponding author on reasonable request.

## Code availability statement

Custom code developed for analyzing the data during the current study is available on: https://github.com/drguggiana/IVIV_pipeline

## Author Contribution

S.W. and V.S. conceived the project, with input from M.H., T.R., and T.B.. S.W. planned and performed all experiments. D.G.N. and S.W. wrote advanced analysis tools and D.G.N., S.W., T.R. and V.S. analyzed the data. S.W. and V.S. implemented LSPS at the patch-clamp setups. T.R. designed and built the *in vivo* 2-photon setup and developed the viral construct. S.W., D.G.N, M.H., V.S., T.R. and T.B. wrote the manuscript. T.B. provided the research environment.

## References

Adolfsson A, Ackerman M, Brownstein NC (2019) To cluster, or not to cluster: An analysis of clusterability methods. Pattern Recognit 88:13–26.

Andermann ML, Kerlin AM, Roumis DK, Glickfeld LL, Reid RC (2011) Functional specialization of mouse higher visual cortical areas. Neuron 72:1025–1039.

Bekkers JM, Häusser M (2007) Targeted dendrotomy reveals active and passive contributions of the dendritic tree to synaptic integration and neuronal output. Proc Natl Acad Sci U S A 104:11447–11452.

Benjamini Y, Hochberg Y (1995) Controlling the False Discovery Rate: A Practical and Powerful Approach to Multiple Testing. J R Stat Soc Ser B 57:289–300.

Bielza C, Benavides-Piccione R, López-Cruz P, Larrañaga P, Defelipe J (2014) Branching angles of pyramidal cell dendrites follow common geometrical design principles in different cortical areas. Sci Rep 4:1–7.

Brainard DH (1997) The Psychophysics Toolbox. Spat Vis 10:433–436.

Callaway EM, Katz LC (1993) Photostimulation using caged glutamate reveals functional circuitry in living brain slices. Proc Natl Acad Sci U S A 90:7661–7665.

Carandini M, Ferster D (2000) Membrane potential and firing rate in cat primary visual cortex. J Neurosci 20:470–484.

Cuntz H, Forstner F, Borst A, Häusser M (2011) The TREES toolbox-probing the basis of axonal and dendritic branching. Neuroinformatics 9:91–96.

Dantzker JL, Callaway EM (2000) Laminar sources of synaptic input to cortical inhibitory interneurons and pyramidal neurons. Nat Neurosci 3:701–707.

Deitcher Y, Eyal G, Kanari L, Verhoog MB, Atenekeng Kahou GA, Mansvelder HD, de Kock CPJ, Segev I (2017) Comprehensive Morpho-Electrotonic Analysis Shows 2 Distinct Classes of L2 and L3 Pyramidal Neurons in Human Temporal Cortex. Cereb Cortex 27:5398–5414.

Gouwens NW et al. (2019) Classification of electrophysiological and morphological neuron types in the mouse visual cortex. Nat Neurosci 22:1182–1195.

Günay C, Edgerton JR, Li S, Sangrey T, Prinz AA, Jaeger D (2009) Database analysis of simulated and recorded electrophysiological datasets with PANDORA’s toolbox. Neuroinformatics 7:93–111.

Gur M, Snodderly DM (2008) Physiological differences between neurons in layer 2 and layer 3 of primary visual cortex (V1) of alert macaque monkeys. J Physiol 586:2293–2306.

Harris KD, Mrsic-Flogel TD (2013) Cortical connectivity and sensory coding. Nature 503:51–58.

Harris KD, Shepherd GMG (2015) The neocortical circuit: Themes and variations. Nat Neurosci 18:170–181.

Hartigan JA (1985) The dip test of unimodality. Ann Stat 13:70–84.

Jiang X, Shen S, Cadwell CR, Berens P, Sinz F, Ecker AS, Patel S, Tolias AS (2015) Principles of connectivity among morphologically defined cell types in adult neocortex. Science (80-) 350:6264.

Kanari L, Ramaswamy S, Shi Y, Morand S, Meystre J, Perin R, Abdellah M, Wang Y, Hess K, Markram H (2019) Objective Morphological Classification of Neocortical Pyramidal Cells. Cereb Cortex 29:1719–1735.

Kätzel D, Zemelman B V., Buetfering C, Wölfel M, Miesenböck G (2011) The columnar and laminar organization of inhibitory connections to neocortical excitatory cells. Nat Neurosci 14:100–109.

Kerlin AM, Andermann ML, Berezovskii VK, Reid RC (2010) Broadly Tuned Response Properties of Diverse Inhibitory Neuron Subtypes in Mouse Visual Cortex. Neuron 67:858–871.

Kim EJ, Juavinett AL, Kyubwa EM, Jacobs MW, Callaway EM (2015) Three Types of Cortical Layer 5 Neurons That Differ in Brain-wide Connectivity and Function. Neuron 88:1253–1267.

Kim EJ, Zhang Z, Huang L, Ito-Cole T, Jacobs MW, Juavinett AL, Senturk G, Hu M, Ku M, Ecker JR, Callaway EM (2020) Extraction of Distinct Neuronal Cell Types from within a Genetically Continuous Population. Neuron 107:274–282.e6.

Kim MH, Znamenskiy P, Iacaruso MF, Mrsic-Flogel TD (2018) Segregated Subnetworks of Intracortical Projection Neurons in Primary Visual Cortex. Neuron 100:1313–1321.e6.

Kreile AK, Bonhoeffer T, Hübener M (2011) Altered visual experience induces instructive changes of orientation preference in mouse visual cortex. J Neurosci 31:13911–13920.

Larkman A, Mason A (1990) Correlations between morphology and electrophysiology of pyramidal neurons in slices of rat visual cortex. I. Establishment of cell classes. J Neurosci 10:1407–1414.

Leinweber M, Zmarz P, Buchmann P, Argast P, Hübener M, Bonhoeffer T, Keller GB (2014) Two-photon calcium imaging in mice navigating a virtual reality environment. J Vis Exp 84:1–6.

Luo H, Hasegawa K, Liu M, Song W-J (2017) Comparison of the Upper Marginal Neurons of Cortical Layer 2 with Layer 2/3 Pyramidal Neurons in Mouse Temporal Cortex. Front Neuroanat 11:115.

Mainen ZF, Sejnowski TJ (1996) Influence of dendritic structure on firing pattern in model neocortical neurons. Nature 382:363–366.

Marshel JH, Garrett ME, Nauhaus I, Callaway EM (2011) Functional specialization of seven mouse visual cortical areas. Neuron 72:1040–1054.

McInnes L, Healy J, Melville J (2018) UMAP: Uniform Manifold Approximation and Projection for Dimension Reduction.

Meng X, Winkowski DE, Kao JPY, Kanold PO (2017) Sublaminar Subdivision of Mouse Auditory Cortex Layer 2/3 Based on Functional Translaminar Connections. J Neurosci 37:10200–10214.

Morgenstern NA, Bourg J, Petreanu L (2016) Multilaminar networks of cortical neurons integrate common inputs from sensory thalamus.

Niell CM, Stryker MP (2008) Highly selective receptive fields in mouse visual cortex. J Neurosci 28:7520–7536.

O’Herron P, Levy M, Woodward JJ, Kara P (2020) An Unexpected Dependence of Cortical Depth in Shaping Neural Responsiveness and Selectivity in Mouse Visual Cortex. eneuro:ENEURO.0497-19.2020.

Peng H et al. (2021) Morphological diversity of single neurons in molecularly defined cell types. Nat 2021 5987879 598:174–181.

Petersen CCH, Crochet S (2013) Synaptic Computation and Sensory Processing in Neocortical Layer 2/3. Neuron 78:28–48.

Petreanu L, Mao T, Sternson SM, Svoboda K (2009) The subcellular organization of neocortical excitatory connections. Nature 457:1142–1145.

Pologruto TA, Sabatini BL, Svoboda K (2003) ScanImage: Flexible software for operating laser scanning microscopes. Biomed Eng Online 2:13.

Rose T, Jaepel J, Hübener M, Bonhoeffer T (2016) Cell-specific restoration of stimulus preference after monocular deprivation in the visual cortex. Science (80-) 352:1319–1322.

Rossi LF, Harris KD, Carandini M (2020) Spatial connectivity matches direction selectivity in visual cortex. Nature 588:648–652.

Scala F, Kobak D, Bernabucci M, Bernaerts Y, Cadwell CR, Castro JR, Hartmanis L, Jiang X, Laturnus S, Miranda E, Mulherkar S, Tan ZH, Yao Z, Zeng H, Sandberg R, Berens P, Tolias AS (2021) Phenotypic variation of transcriptomic cell types in mouse motor cortex. Nat 2020 5987879 598:144–150.

Schindelin J, Arganda-Carreras I, Frise E, Kaynig V, Longair M, Pietzsch T, Preibisch S, Rueden C, Saalfeld S, Schmid B, Tinevez JY, White DJ, Hartenstein V, Eliceiri K, Tomancak P, Cardona A (2012) Fiji: An open-source platform for biological-image analysis. Nat Methods 9:676–682.

Shepherd GMG, Stepanyants A, Bureau I, Chklovskii D, Svoboda K (2005) Geometric and functional organization of cortical circuits. Nat Neurosci 8:782–790.

Shepherd GMG, Svoboda K (2005) Laminar and columnar organization of ascending excitatory projections to layer 2/3 pyramidal neurons in rat barrel cortex. J Neurosci 25:5670–5679.

Staiger JF, Bojak I, Miceli S, Schubert D (2015) A gradual depth-dependent change in connectivity features of supragranular pyramidal cells in rat barrel cortex. Brain Struct Funct 220:1317–1337.

Suter BA, O’Connor T, Iyer V, Petreanu LT, Hooks BM, Kiritani T, Svoboda K, Shepherd GMG (2010) Ephus: Multipurpose data acquisition software for neuroscience experiments. Front Neural Circuits 4:1–12.

Tasic B et al. (2016) Adult mouse cortical cell taxonomy revealed by single cell transcriptomics. Nat Neurosci 19:335–346.

Tasic B et al. (2018) Shared and distinct transcriptomic cell types across neocortical areas. Nature 563:72–78.

Tyler WA, Medalla M, Guillamon-Vivancos T, Luebke JI, Haydar TF (2015) Neural precursor lineages specify distinct neocortical pyramidal neuron types. J Neurosci 35:6142–6152.

Van Aerde KI, Feldmeyer D (2015) Morphological and physiological characterization of pyramidal neuron subtypes in rat medial prefrontal cortex. Cereb Cortex 25:788–805.

Vélez-Fort M, Rousseau C V., Niedworok CJ, Wickersham IR, Rancz EA, Brown APY, Strom M, Margrie TW (2014) The stimulus selectivity and connectivity of layer six principal cells reveals cortical microcircuits underlying visual processing. Neuron 83:1431–1443.

Weiler S, Bauer J, Hübener M, Bonhoeffer T, Rose T, Scheuss V (2018) High-yield in vitro recordings from neurons functionally characterized in vivo. Nat Protoc 13:1275–1293.

Weiler S, Nilo DG, Bonhoeffer T, Hübener M, Rose T, Scheuss V (2020) Relationship between input connectivity, morphology and orientation tuning of layer 2/3 pyramidal cells in mouse visual cortex. bioRxiv:2020.06.03.127191.

Xu X, Olivas ND, Ikrar T, Peng T, Holmes TC, Nie Q, Shi Y (2016) Primary visual cortex shows laminar-specific and balanced circuit organization of excitatory and inhibitory synaptic connectivity. J Physiol 594:1891–1910.

Yao Z et al. (2021) A taxonomy of transcriptomic cell types across the isocortex and hippocampal formation. Cell 184:1–20.

Zaitsev A V., Povysheva N V., Gonzalez-Burgos G, Lewis DA (2012) Electrophysiological classes of layer 2/3 pyramidal cells in monkey prefrontal cortex. J Neurophysiol 108:595–609.

